# Morphological evolution in the ant reproductive caste

**DOI:** 10.1101/2020.07.18.210302

**Authors:** Raquel Divieso, Thiago S. R. Silva, Marcio R. Pie

**Affiliations:** Departamento de Zoologia, Universidade Federal do Paraná, CEP 81531-990, Curitiba, Paraná, Brazil

**Keywords:** males, queens, sexual dimorphism, Rensch’s rule

## Abstract

The evolution of eusociality led to severe changes in the general hymenopteran body plan. In particular, the evolution of reproductive division of labour caused the worker caste to be largely freed from the demands involved in reproduction. As a consequence, workers were able to evolve highly specialized morphologies for foraging and colony maintenance, whereas the reproductive caste became specialized for reproduction. Despite these important changes, little is known about general patterns of morphological evolution within the ant reproductive caste. Our goals were to characterize morphological variation in the ant reproductive caste and to test whether different sexes display variation in their evolutionary rates. We obtained measurements of 897 specimens from a total of 678 ant species. The shapes of the size distributions were similar between sexes, with queens being larger than males in all traits except for eye length. Contrary to the expectation based on Rensch’s rule, although queens were larger, the degree of dimorphism increased with body size. Finally, there is strong evidence for an accelerated tempo of morphological evolution in queens in relation to males. These results represent the first comprehensive treatment of morphological variation in the ant reproductive caste and provide important new insights into their evolution.

## INTRODUCTION

Eusociality is traditionally defined by the presence of three main characteristics: cooperative care of young, overlapping generations, and reproductive division of labour (Wilson 1971, Wilson & Hölldobler, 2005). Although the first two characteristics are often found in many organisms with relatively low levels of sociality, as in the case of cooperative breeding in birds (Koenig & Dickinson, 2004) and mammals (Clutton-Brock, 2006), reproductive division of labour is both phylogenetically uncommon and most challenging to explain, ever since it was first framed as Darwin’s “special difficulty” for the theory of evolution by natural selection in the Origin of Species. Although the conundrum of the evolution of reproductive division of labour has been assuaged by the advent of kin selection theory (Hamilton, 1964a,b), its implications for the morphological evolution of the reproductive caste are still poorly understood. One of the consequences of reproductive division of labour, particularly in highly eusocial species, is that the worker caste becomes largely released from the demands involved with reproduction and therefore could evolve highly specialized morphologies for foraging and colony maintenance, whereas the reproductive caste might invest more heavily on reproduction (Hölldobler & Wilson 1990). As a consequence, the reproductive caste effectively functions as the gonads of a superorganism (colony), possibly being able to evolve a level of morphological specialization that would otherwise be unattainable in a solitary insect.

Another important consequence of the evolution of the reproductive caste involves the differences between males and queens in haplodiploid species (Stubblefield & Seger, 1994; Paxton, 2005). In general, males do not take part in colony tasks and are expected to be short-lived once they leave the nest (Boomsma *et al*., 2005; however, see Shik *et al*. [2013] for a discussion on the male life-history continuum hypothesis). On the other hand, queens are responsible for colony foundation and often show extraordinary lifespans (Keller & Genoud, 1997). These asymmetries are likely to have a profound impact on the selective pressures for different morphological traits within and among sexes, as well as in their rates, yet little is known about general trends in morphological evolution of ant males and queens. Most research on ant male evolution has focused on mating, reproductive investment, and variation in reproductive organs (e.g. Baer & Boomsma, 2004; Boomsma *et al*., 2005; Baer 2011; Boudinot, 2013). On the other hand, variation in thorax morphology between queens and workers is associated with the functions performed by the worker caste, with nesting strategies accentuating these differences (Keller *et al*., 2014; Peeters *et al*., 2020 unpubl. data), yet the relative scarcity of comparable information on males currently prevents large-scale comparisons in traits, particularly those not involved in mating.

In this study, we investigate the tempo and mode of morphological evolution in the ant reproductive caste. In particular, our goals were: (1) to characterize morphological variation in the ant reproductive caste; (2) to assess the level of sexual dimorphism; (3) to estimate the degree of phylogenetic signal in male and queen traits; and (4) to test whether different sexes display differences in their rates of evolution.

## MATERIAL AND METHODS

High-resolution images of 897 specimens (552 queens and 345 males – lateral, dorsal, and full-face views) were obtained for a total of 678 ant species using the AntWeb platform (http://www.antweb.org). Whenever possible, we preferably selected specimens corresponding to holotypes or paratypes. We did not include ergatoid males, such as those found in *Cardiocondyla* (e.g. Heinze *et al*., 2005), because they represent a qualitatively different type of reproductive strategy than winged males. Species names, raw measurements, and corresponding voucher numbers are indicated in Table S1. Six measurements were obtained from each specimen (Figure 1): eye length (EL), distance between the eyes (BE), head width (HW), head length (HL), mesosoma length (ML), and mesonotum width (MW). Scale bars on each image were used for calibration. Some measurements were missing (eight out of 6286 measurements) because the position of the specimen prevented its reliable observation. All measurements were obtained using ImageJ 1.49 (Schneider *et al*., 2012).

**Figure 1.**
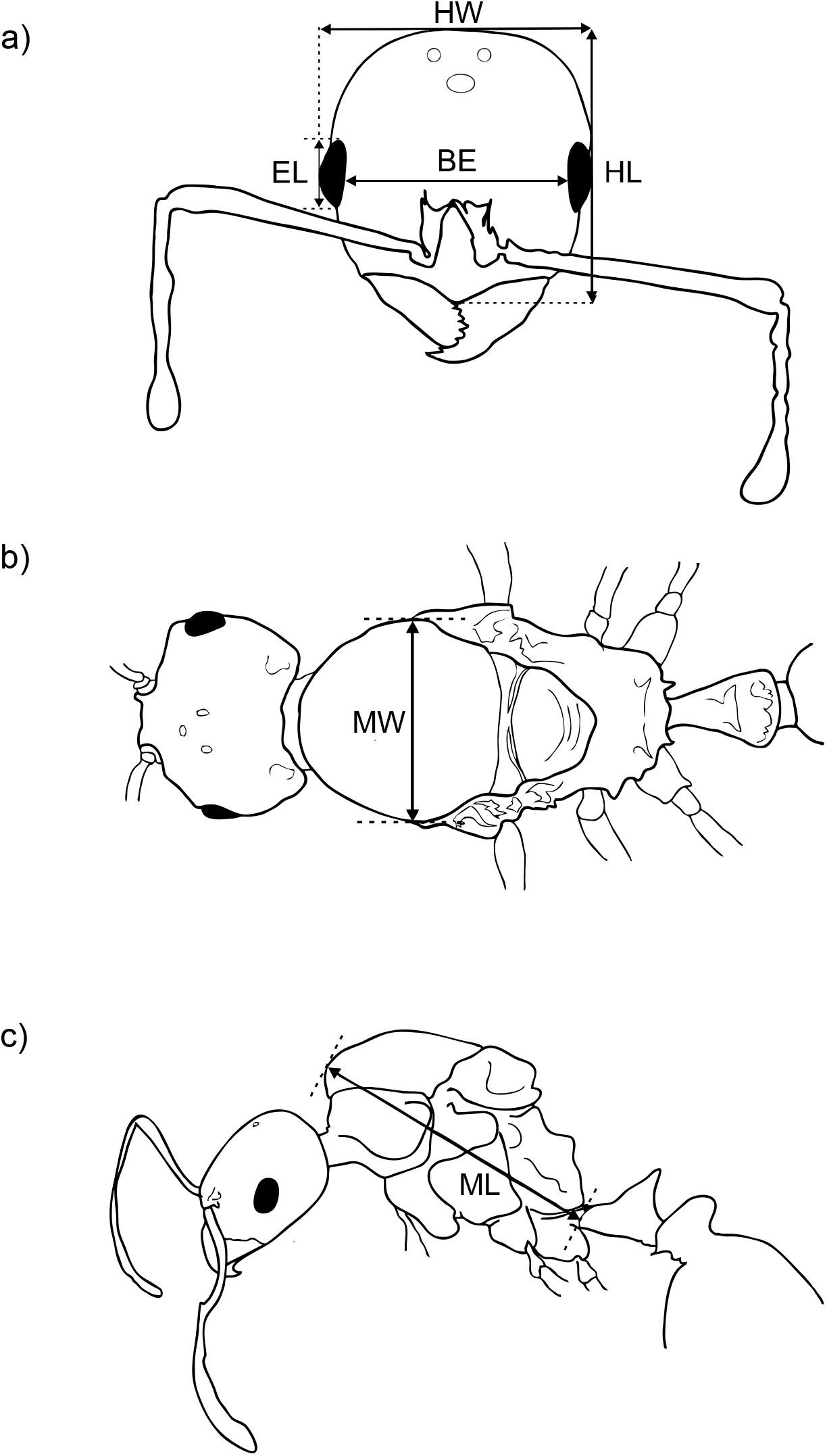
Morphological measurements used in the present study. Measurements in full-face view: head width (HW), head length (HL), eye length (EL), distance between the eyes (BE), in lateral view: mesosoma length (ML); in dorsal view: mesonotum width (MW).

We began our morphological analyses by exploring the distribution of each measured trait in males and queens, followed by an assessment of the degree of sexual dimorphism by plotting male and queen traits against one another, given the expectation that the absence of dimorphism would lead to a relationship with a slope of 1. We also used the index of Lovich & Gibbons (1992), calculated as the (size of the queen trait/size of the male trait) – 1. The main advantage of this index is that it is easily interpretable, with positive values indicating relatively larger queens and negative values indicating relatively larger males. In order to explore how dimorphism evolved over the course of ant evolution, we mapped indices for different traits onto the ant phylogeny (see below) using the contMap function in the phytools 0.6-44 package (Revell, 2012). Sexual dimorphism analyses involved a subset of 218 species for which male and queen data were available.

We then explored morphological variation in the reproductive caste using principal component analyses (PCAs) based on covariance matrices of log-transformed measurements using the function prcomp in STATS 3.4.2 in R (R Core Team, 2020). In a preliminary PCA, PC1 accounted for more than 90% of the variation in the dataset. This is clearly due to the strong pairwise correlation among all measured traits (Figure S1) indicating a strong influence of body size which could mask important morphological variation. Therefore, we used the regression residues of each trait against PC1 in a new PCA analysis to decrease the effect of size on the remaining morphological variation. PCAs were carried out for each sex separately. PCs retained for further interpretation were selected using the broken-stick method, which considers as interpretable components those with observed eigenvalues higher than the variance produced by the broken stick method (Jackson 1993).

We assessed the degree of phylogenetic signal of each trait in males and queens using the λ parameter of Pagel (1999). More specifically, we test whether a model with an estimated λ provides a better fit than a simpler alternative where λ=0. Parameter estimates were obtained using the phytools 0.6-44 package (Revell, 2012). We estimated the rate of evolution of each trait based on the σ^2^ parameter using the fltContinuous function in geiger 2.0.7 (Harmon *et al*., 2008) under the corresponding estimated value of λ for each clade. Differences in rates of evolution between male and queen traits were tested using the mvBM function in mvMORPH 1.1.3 (Clavel *et al*., 2015). This test contrasts the fit of a model in which males and queens share the same rates against an alternative where each sex has a separate rate using their AIC scores.

Given that there is no available species-level phylogeny for the studied species, we used a two-step process to incorporate phylogenetic information in our estimates of phylogenetic signal and evolutionary rates. We began by building a composite phylogeny based on the tree by Moreau & Bell (2013), but using the phylogenetic relationships and divergence times within Myrmicinae from Ward *et al*. (2015). This phylogeny was then pruned to keep only a single species per genus to generate a genus-level phylogeny. We then used the list of species in our dataset to simulate a species-level phylogeny in which the relationships within genera were obtained from a Yule (pure-birth) process using the genus.to.species.tree function in phytools. A total of 500 simulated trees were used in downstream analyses. All analyses were carried out in R 4.0.2 (R Core Team, 2020).

## RESULTS

The shapes of the distributions of all measured traits tended to be similar, with queens being larger in all traits, except for eye length (EL, Figure 2, Table S2). However, when traits were plotted against each other, there was a clear pattern of sexual dimorphism, which tended to increase with body size (Figure 3). In particular, dimorphism was most commonly biased toward larger queens in all traits, except for EL. Interestingly, when dimorphism for each trait was mapped onto the phylogeny, a remarkably complex pattern emerged, with the degree and direction of dimorphism varying considerably among traits (Figure 4). In general, the ancestral condition in ants seems involve some moderate level of dimorphism involving larger queens, which became more pronounced independently in several lineages for HW, HL, BE and ML, as well as a few cases of the dimorphism in the other direction, with larger males. On the other hand, MW showed a more complex pattern, with several independent instances of the evolution of disproportionately larger males. Finally, a qualitatively different pattern is found in EL, with negative dimorphism levels (relatively larger traits in males) in the poneroid clade giving rise to moderately positive values at the base of the formicoid clade and finally reverting back to negative dimorphism in myrmicines.

**Figure 2.**
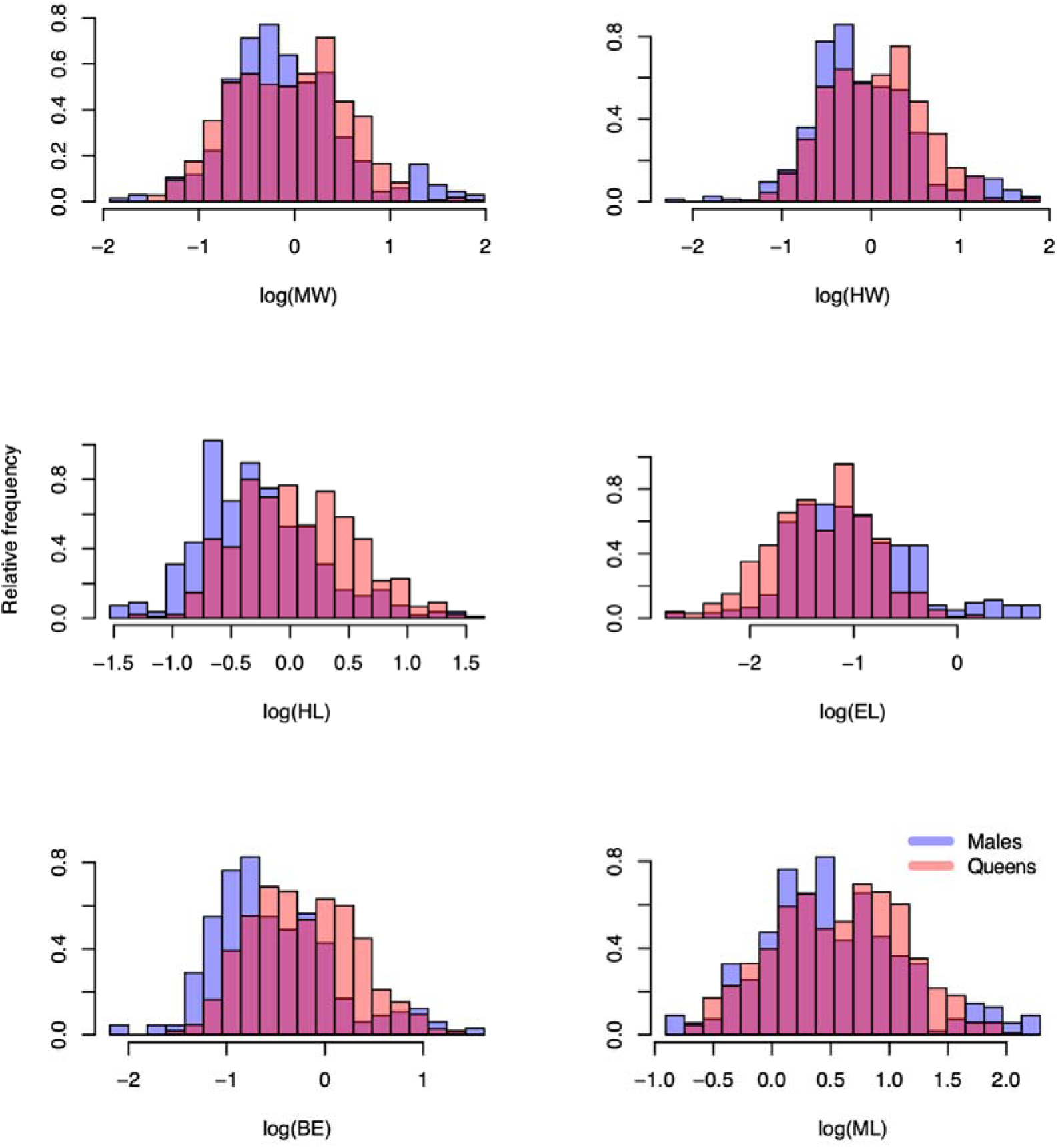
Relative distribution of measurements of males and queens across the studied species: head width (HW), head length (HL), eye length (EL), distance between the eyes (BE), mesosoma length (ML), mesonotum width (MW).

**Figure 3.**
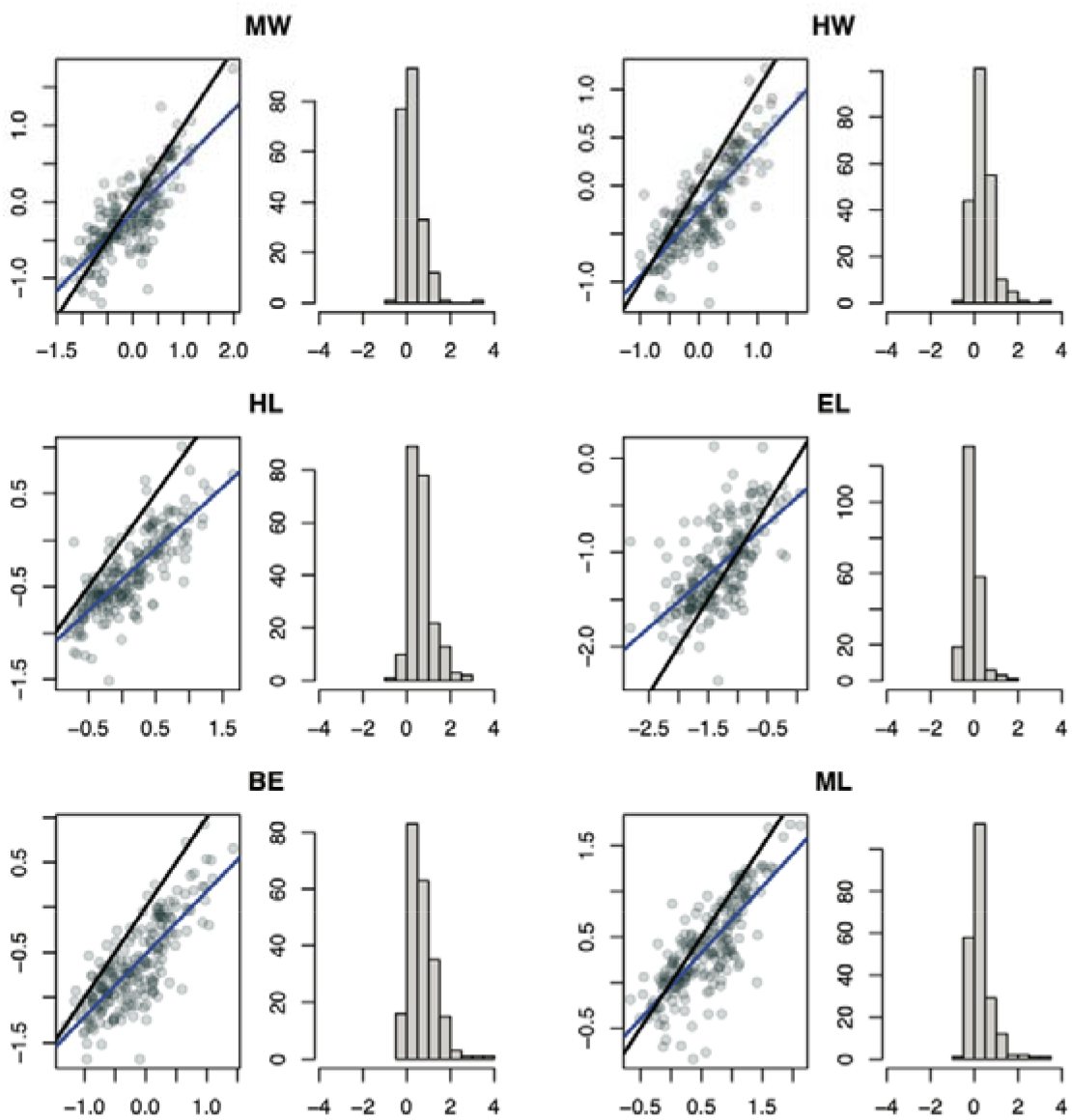
Relationship between male (x axis) and female (y axis) traits in the ant reproductive caste and the corresponding level of sexual dimorphism based on the index of Lovich & Gibbons (1992) : head width (HW), head length (HL), eye length (EL), distance between the eyes (BE), mesosoma length (ML), mesonotum width (MW).

**Figure 4.**
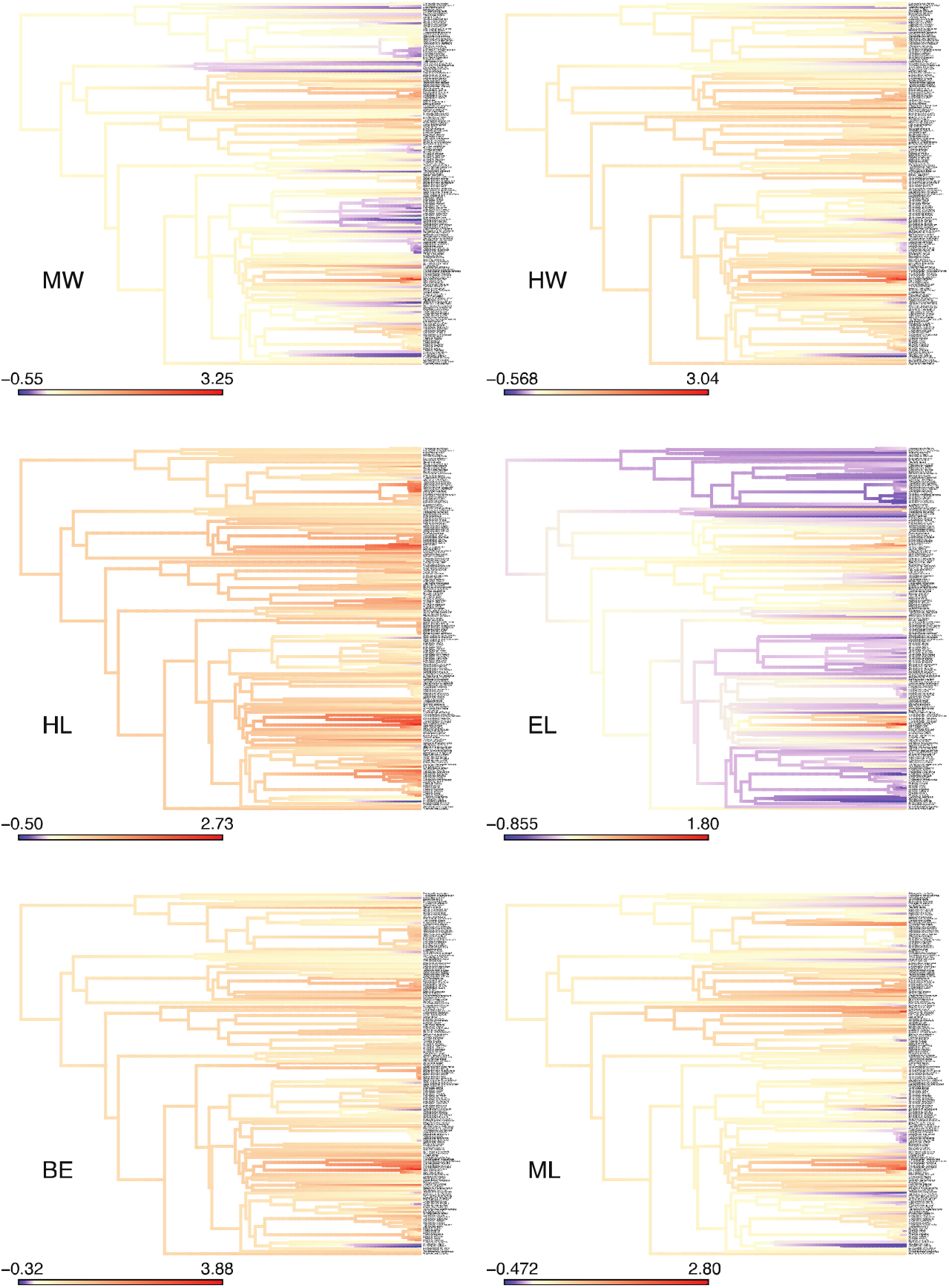
Ancestral state reconstruction of the evolution of sexual dimorphism in ants based on the index of Lovich & Gibbons (1992). The used tree was randomly chosen among the set of alternative topologies (see text for details), but the resulting reconstructions were largely unaffected by differences between trees.

PCA loadings can be found on Table 1. The first PC explained 44% of the total variance in queens and 36% in males, while PCs 2 and 3 explained 24% and 17% in both sexes. The first PC in queens mostly reflected overall head width, with the highest loadings being found on HW and BE. PC2 and PC3 in queens described negative relationships between MW x HL and EL x ML, respectively. Males showed a qualitatively different organization in their PCA. Positive scores on PC1 described more elongate males, with narrow heads and long mesosomas (Table 1). PC2 in males reflected a different trade-off, with positive scores being associated with eyes that are both large and distant from one another irrespective of head width. Finally, PC3 in males mostly reflected HL.

**Table 1.** Loadings of the principal component analyses of morphological variation in ant males and queens. Males and queens were analysed separately using log-transformed data.

| **Traits** | **Queens** |  |  | **Males** |  |  |
| --- | --- | --- | --- | --- | --- | --- |
|  | **PC1** | **PC2** | **PC3** | **PC1** | **PC2** | **PC3** |
| MW | -0.24 | 0.69 | -0.17 | 0.47 | 0.35 | -0.23 |
| HW | 0.54 | 0.12 | 0.17 | -0.50 | 0.18 | -0.39 |
| HL | 0.30 | -0.60 | -0.13 | -0.31 | -0.23 | 0.74 |
| EL | -0.37 | -0.21 | 0.73 | -0.18 | -0.67 | -0.39 |
| BE | 0.54 | 0.17 | -0.09 | -0.35 | 0.57 | 0.17 |
| ML | -0.36 | -0.29 | -0.63 | 0.52 | -0.13 | 0.25 |
| % explained variance | 44% | 25% | 17% | 36% | 25% | 17% |

Phylogenetic signal was very strong in all measured traits for both sexes (Figure S2), with λ values close to one regardless of variation in the used tree topologies (p< 3.84e-10 in all tests). On the other hand, there were some intriguing differences when phylogenetic signal was tested on the PC scores for each sex. Males showed lower but still significant phylogenetic signal on PC2, whereas the phylogenetic signal for males on this axis was still high (lowest P value across all 1000 alternative trees = 0.0145 and 2.15e-22, respectively). Finally, a much lower phylogenetic signal was marginal on PC3 for both sexes, with P values between 0.00007 and 0.30 depending on the topology.

There were substantial differences in rates of evolution in all measured traits (Figure 5). The difference was more pronounced when traits were tested separately, as it tends to conflate the effects of body size and shape, although variation along the size-corrected PCA also showed differences. In all comparisons, the model with separate rate parameters showed on average ΔAIC between 35.8 and 56.6, which suggests overwhelming support for a difference in rates between sexes. In particular, queens evolved faster than males in all traits, except for PC3. The latter difference is intriguing given that PC3 was mostly associated with HW, which evolved faster in males. However, it is important to note that we used size-corrected residuals for the PCA, so that, even though absolute values of HL indeed evolved faster on queens, males evolved HL faster than queens in proportion to their body size.

**Figure 5.**
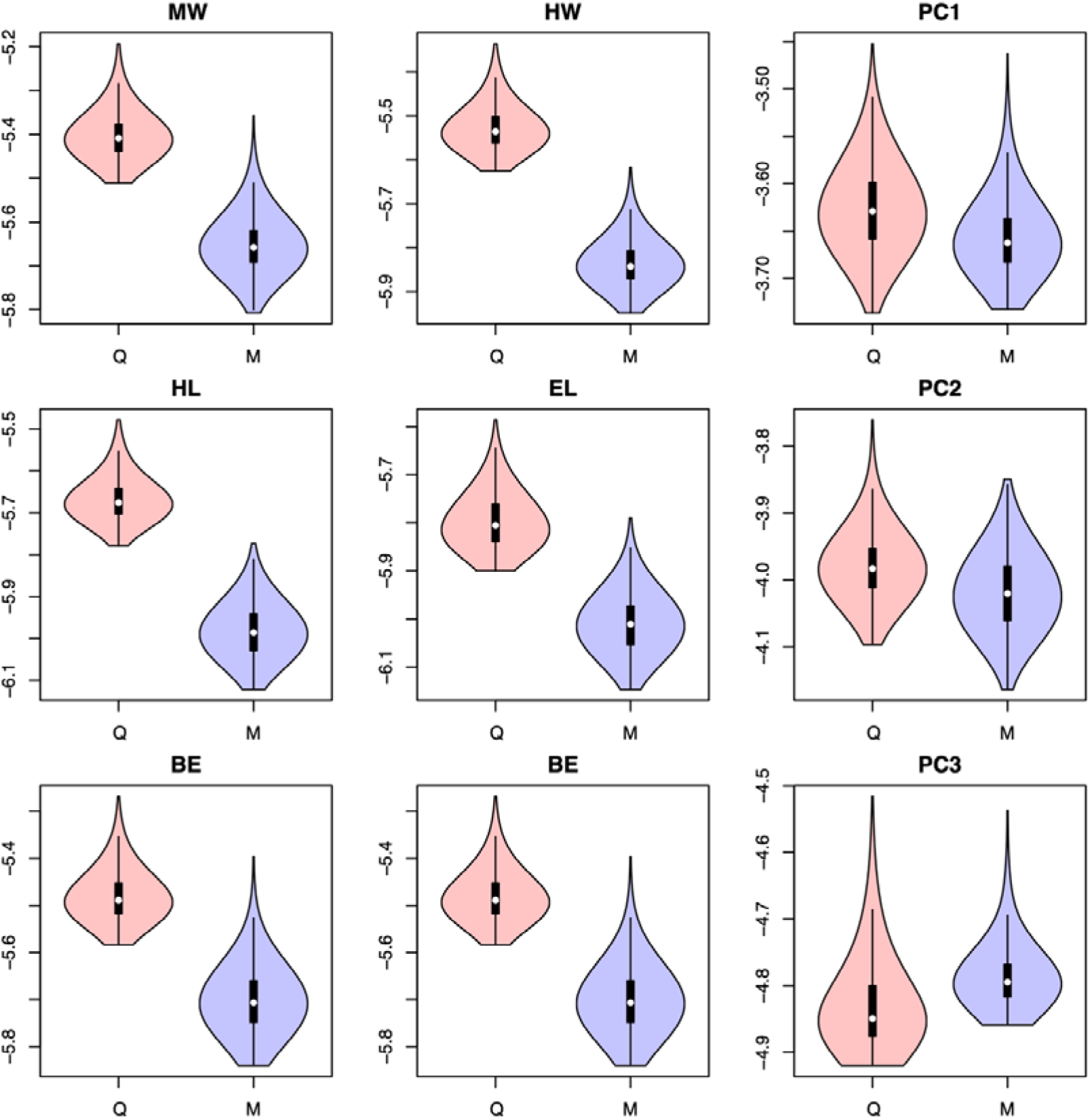
Variation in evolutionary rates between queens and males for each measured trait, as well as the scores on the first principal component axes. See text for details.

## DISCUSSION

Despite the substantial morphological variation between and within males and females in the ant reproductive caste uncovered in our analyses, we identified several robust patterns governing their evolution, namely (1) there was substantial sexual dimorphism in ants, with queens being larger than males and the degree of dimorphism increasing with body size, except for eye length (EL); (2) different body regions had qualitatively different evolutionary trends, underscoring the modular nature of their body plan; and (3) the tempo of morphological evolution was faster in queens for all traits, except for EL. Each of these principles will be discussed in turn below.

According to Rensch’s rule, size dimorphism should decrease with increasing average body size when the queens is the larger sex (Rensch, 1950). Although the existence of Rensch’s rule has never been tested for ants before, evidence from previous studies suggest that it is uncommon among insects (Teder & Tamaru, 2005; Blanckenhorn *et al*., 2007a). Indeed, our study found evidence for the opposite of Rensch’s rule, with the level of dimorphism increasing with body size despite queens being larger (Figure 3). A similar discrepancy, but in a smaller magnitude, in relation to the prediction of Rensch’s rule was found in solitary wasps (Blanckenhorn *et al*., 2007b; Fairbairn *et al*., 2007), but other hymenopterans found either no allometry (“primitive” bees [Del Castillo & Fairbairn, 2012]) or conflicting results (Meliponini bees [Quezada-Euán *et al*., 2019]). However, it is important to note that these large-scale patterns found in our study belie a remarkable variation in the degree of dimorphism among ant lineages (Figure 4) and emphasize how little is known about the life-history and sexual selection mechanisms driving such variation in dimorphism.

The body in the ant reproductive caste plan of ants has been interpreted as inherently modular (Molet *et al*., 2012), with different body regions responding to specific demands, particularly with respect to colony foundation in the case of queens (Keller *et al*., 2014), although little is known about this with respect in males. Such modular organization seems to extend to differences among sexes as well. For instance, the differences found in values and associated rates for HW and HL, for example, provide hints of differential functional and anatomical constraints acting on the evolution of traits between sexes. In the case of cephalic size variation, most of the internal surface of the ant head act as support site for the muscles that attach to the mouthparts and antennae (Richter *et al*., 2019; 2020). This is particular true for the mandibular muscles (i.e. anterior cranio-mandibular and posterior cranio-mandibular muscles [0md1 and 0md3 in Richter *et al*., 2020, respectively]), which occupy much of the space of the internal surface of the head capsule in workers (Richter *et al*., 2020). Variation of cephalic values between queens and males are possibly constrained by differential mandibular usage during life history. Despite the relaxation of mandibular use by queens after founding (especially in species with claustral strategies), they retain functional importance in different periods of its life history. In males, however, most groups have simplified of mouthparts (especially mandibles), with the putative reduction of muscle volume and area of attachment, reflecting the sub-usage those components in the life history of this particular sex (Gronenberg *et al*., 1997), releasing them from the functional constraints that determine size variation in the head.

Considering the variation of the MW and its putative constraints, we have to consider that the ant mesonotum has an intimate relationship with the head, given that it internally corresponds to the site of origin of the pronoto-laterocervical, pronoto-postoccipital and pronoto-propleural muscles (Keller *et al*., 2014; Silva & Feitosa, 2019), which are mainly responsible for most movements of the head capsule in hymenopterans (Mikó *et al*., 2007; Snodgrass, 1942). According to Moll *et al*. (2010), precise head movements are essential to reduce displacement of the centre of mass, and retain stability while carrying objects many times the individuals’ weight and length, with the pronotal size variation (and related skeletomuscular modifications) playing a major role in achieving this biomechanical function (see also Anderson *et al*., 2020 and Keller *et al*., 2014). In queens, mesonotum width (MW) was most strongly associated with PC2, which was dominated by a negative relationship between mesonotum width and head length (HL). On the other hand, MW in males was most strongly associated with PC1, varying in the same direction as mesosoma length (ML) and negatively with head width (HW). These results suggest that MW may be a variable that better explains morphological variation associated to dispersion strategies rather than cephalic mobility. Pronotal width variation may be constrained by width variation of the prophragma-bearing sclerite (i.e. mesoscutum), which receives the muscular components related to flight (e.g. the prophragmo-mesophragmal muscles) (Vilhelmsem *et al*., 2010). As mentioned by Peeters *et al*. (2020 unpubl. data), the space of the prothoracic region is reduced in winged ants, whilst the mesothoracic space is proportionately larger, providing a broader area for attachment of the indirect flight muscles.

The considerable variation in the direction and magnitude of the correlation structure of males and females is consistent with a considerable reorganization of their development to respond to sex-specific life-history demands. Interestingly, the trait which differs the most among sexes in terms of patterns of variation, dimorphism, and evolutionary trajectory is EL, suggesting very different constraints in eye evolution between males and queens. One possibility is that males have a disproportionately larger role in finding conspecific reproductive queens. According to Narendra *et al*. (2016), optical recognition in males is extremely important for mate location and during courtship. Most males tend to have a higher number of ommatidia (although also being comparatively smaller) if compared to congeneric queens, providing them with a higher sampling resolution, despite overall lower optical sensitivity (Narendra *et al*., 2016).

The remarkable faster rate of evolution in queen traits is far from obvious. Indeed, queens have strong selection pressures, especially during the colonial founding stage. In particular, anatomical variation of the mesosoma in queens appears to be correlated to specific strategies of colonial founding (*i.e*. claustral, non-claustral and dependent founding, see Keller *et al*., 2014) which would not directly affect males. On the other hand, if we were to find the opposite pattern, with faster evolution in males, one could argue that the relatively shorter lifespan and the lack of demands to carry out colony tasks could lead to accelerated rates of evolution. Therefore, at the present our interpretations regarding the causes of such interspecific rate differences are necessarily ad hoc and should be explored in further studies, particularly by explicitly testing for correlates of rate variation, including colony size, climate and social organization.

It is important to note that our dataset was not amenable to assessing intraspecific (and intracolonial) variation in males and queens. In particular, we cannot determine the extent to which the patterns found in our analyses are consistently found within species as well. In addition, despite our large dataset, we sampled only a small fraction of the morphological variation found among ant species, and even for those species, we were only able to measure the images that were available. In addition, measurement errors based on variation in the position of the specimens when photographed probably increased the noise in our measurements. Although these sources of error might decrease the power of our analyses, they are unlikely to positively mislead our conclusions. However, precise measurements, in association with more detailed ecological information about focal ant lineages, might prove to be an invaluable tool to further explore the questions raised here, particularly in the contrast between intra- and interspecific variation patterns (Bolnick *et al*., 2011).

In this study we provide the most comprehensive analysis of morphological evolution in the ant reproductive caste to date. Several obvious patterns, such as the substantial variation in sexual dimorphism, the unique pattern of evolution of eye size between and among sexes, and the intriguing temporal evolution of different ant lineages are obvious patterns that had not even been recognized in previous studies. We hope our analyses will provide a basis for new insights into the about the processes and mechanisms driving morphological evolution in ant males and queens.

## ACKNOWLEDGEMENTS

RD and TSRS were funded through graduate and post-doctoral fellowships from CNPq and CAPES, respectively. We thank Luiz R. R. Faria Jr and Roberto Keller for valuable comments on the manuscript.

---

**Figure S1.**
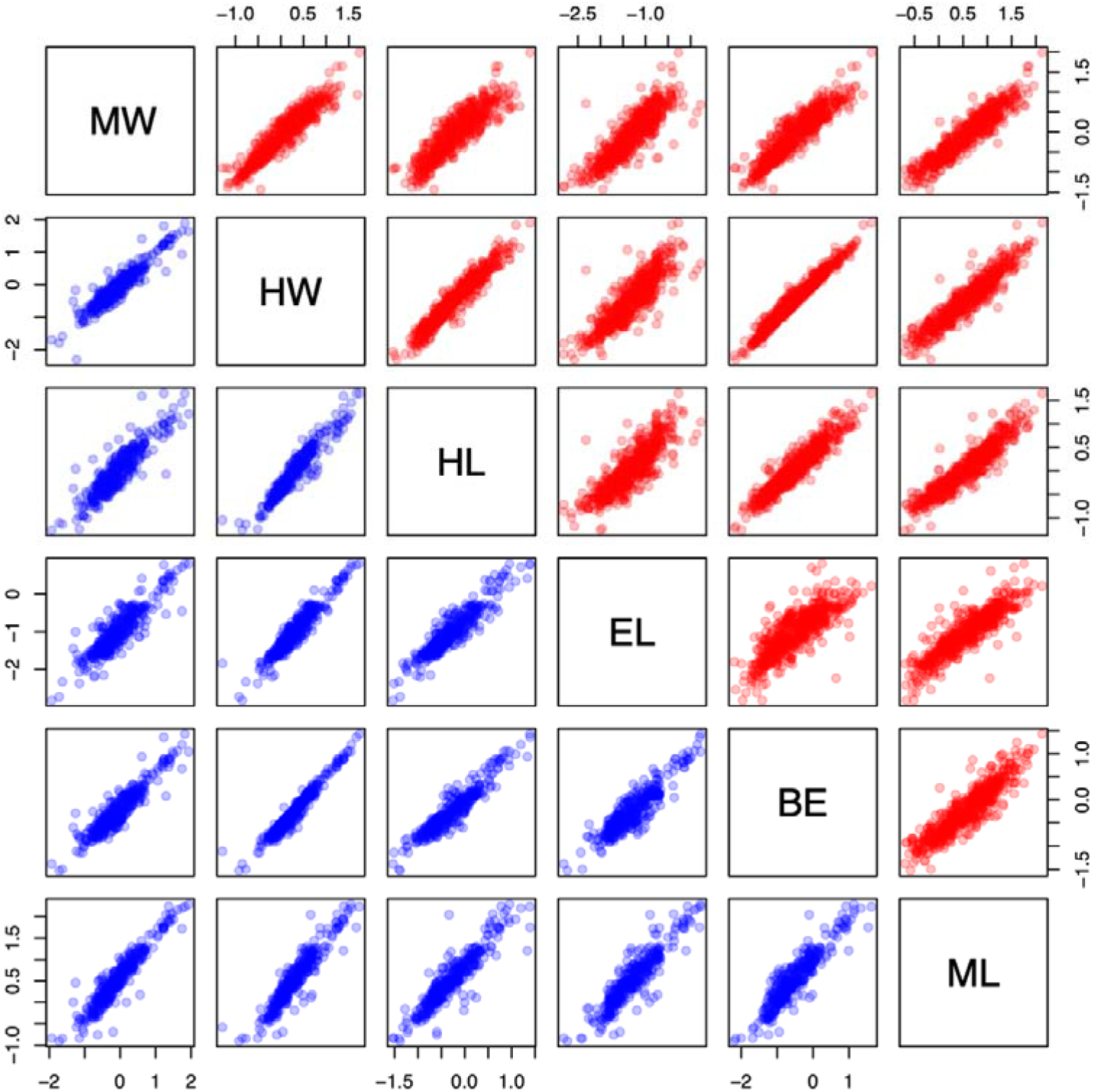
Pairwise scatterplots of queen (red) and male (blue) traits: head width (HW), head length (HL), eye length (EL), distance between the eyes (BE), mesosoma length (ML), mesonotum width (MW).

**Figure S2.**
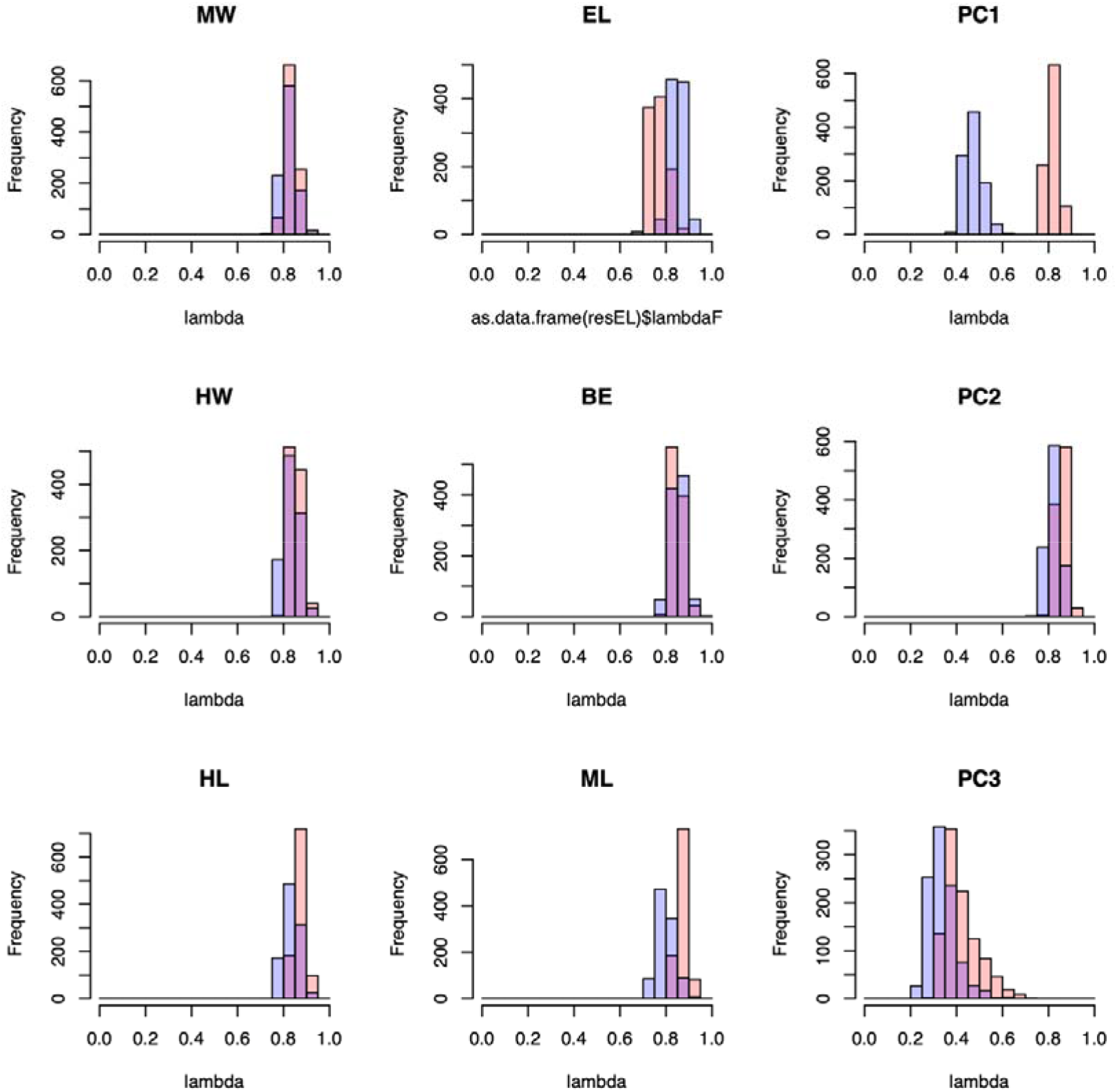
Phylogenetic signal of the measured traits, as well as the first three principal components of their morphological variation. Queens and males are represented by red and blue bars, respectively.

## Supplementary material

Table S1. Species names, raw measurements, and corresponding voucher numbers of the specimens included in this study.

Table S2. Descriptive statistics of the studied variables, presented separately for each ant subfamily. Measurements are indicated as means ± SD (range). N_sp_ and N_gen_ indicates the number of species and genera, respectively.

